# Data-Driven Optimization of DIA Mass Spectrometry by DO-MS

**DOI:** 10.1101/2023.02.02.526809

**Authors:** Georg Wallmann, Andrew Leduc, Nikolai Slavov

## Abstract

Mass spectrometry (MS) enables specific and accurate quantification of proteins with ever increasing throughput and sensitivity. Maximizing this potential of MS requires optimizing data acquisition parameters and performing efficient quality control for large datasets. To facilitate these objectives for data independent acquisition (DIA), we developed a second version of our framework for data-driven optimization of mass spectrometry methods (DO-MS). The DO-MS app v2.0 (do-ms.slavovlab.net) allows to optimize and evaluate results from both label free and multiplexed DIA (plexDIA) and supports optimizations particularly relevant for single-cell proteomics. We demonstrate multiple use cases, including optimization of duty cycle methods, peptide separation, number of survey scans per duty cycle, and quality control of single-cell plexDIA data. DO-MS allows for interactive data display and generation of extensive reports, including publication quality figures, that can be easily shared. The source code is available at: github.com/SlavovLab/DO-MS.

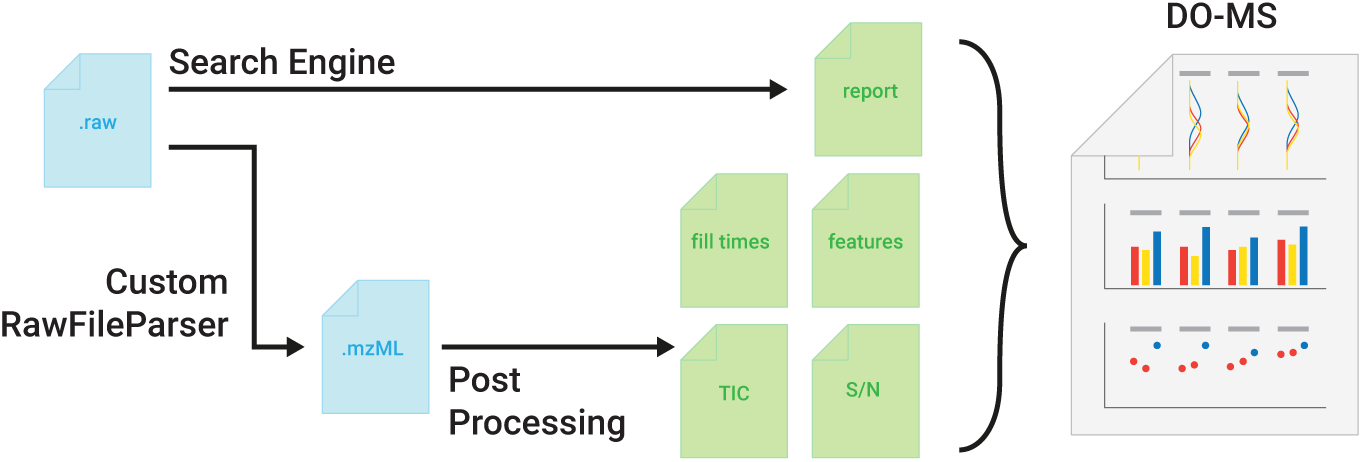

## Introduction

Mass spectrometry (MS) allows for comprehensive quantification and sequence identification of proteins from complex biological samples^1^. Reliable sequence identification of peptides by MS relies on the fragmentation of peptides^2^. This can be performed for one precursor at a time, as in the case of data dependent acquisition (DDA) or for multiple precursors in parallel as in the case of data independent acquisition (DIA). Using real-time instrument control for DDA can achieve high sensitivity, depth and data completeness^3,4^ but remains limited to fragmenting only a subset of the available precursors. This limitation is relaxed by DIA, which systematically selects groups of precursors for fragmentation which cover the whole m/z range^5,6^. This parallel analysis of multiple precursors can have many benefits, including: (1) consistent collection of data from all detectable peptides^7^, (2) high sensitivity due to long ion accumulation times^8^, and (3) high-throughput due to the parallel data acquisition^9^. Despite these benefits, parallel fragmentation of all precursors within the isolation window results in highly complex spectra.

This complexity initially challenged the interpretation of DIA spectra, but advances in machine learning and computational power have gradually increased sequence identification from DIA spectra. Initial approaches were based on sample-specific spectral libraries, but newer methods have allowed for direct library-free DIA and deeper proteome coverage^10–14^. Many current approaches use computationally predicted peptide properties (libraries),^15^ which removes the overhead of experimentally generated libraries. These improvements continue with new acquisition methods^16–18^ and contribute to achieving high proteome depth, data completeness, reproducibility, and throughput^19,20^. This has enabled the quantitative analysis of proteomes down to the singlecell level^21–24^, and can continue to increase the throughput and accuracy of single-cell proteomics towards its biological applications^25^.

Orthogonal to the acquisition method, performance can be further increased when labeling samples with non-isobaric mass tags and analyzing them with the plexDIA framework^26–28^. Multiple labeled samples can be combined and analysed in a single acquisition, multiplicatively increasing the number of protein data points^29^. At the same time, quantitative accuracy and proteome coverage are preserved. As identifications can be translated between different samples labeled by non-isobaric mass tags^26^.

To further empower these emerging capabilities, we sought to extend the DO-MS app to optimization and quality control of DIA experiments by developing and releasing its second major version, v2.0. Indeed, optimization of DIA workflows requires setting multiple acquisition method parameters, such as the number of MS1 survey scans and the placement of fragmentation windows. These parameters must be simultaneously optimized for multiple objectives, including throughput, sensitivity and coverage. Defining the optimal acquisition method therefore becomes a multiobjective, multi-parameter optimization^30,31^. Many tools already exist which cover some aspects of method optimization, like MS2 window placement^18,32,33^. Others focus on quality control^34,35^. DO-MS takes a different approach and offers a holistic view of the acquisition and data processing method specifically designed to diagnose analytical bottlenecks^31^. With this release, DO-MS v2.0 can be used with both DDA data like MaxQuant and DIA data from tools like DIA-NN while having an open interface allowing for adoption to other search engines.

DO-MS is particularly useful for optimizing single-cell proteomic and plexDIA analysis by displaying numerous features relevant to these workflows. These features include intensity distributions for each channel of n-plexDIA^27,29^ and ion accumulation times, which are useful for optimizing single-cell analysis^36,37^, particularly when using isobaric and isotopologous carriers^27,38^. In addition to optimization, DO-MS also facilities data quality control and experimental standardization with large sample cohorts, especially large scale single-cell proteomic experiments^39,40^. Here we demonstrated how DO-MS helps achieve these aims in concrete use cases.

## Results

We developed DIA-specific modules of the DO-MS app^31^ to enable monitoring and optimization of DIA experiments. The DO-MS v2.0 app consists of two parts: A post-processing step which collects additional metrics on the performance of the acquisition method in use and an interactive application to visualize the metrics and results reported by DIA search engines, Fig. 1. All components are built in a modular way, which allows creating new visualization modules and extending the input source to other search engines (the default engine is DIA-NN^13^). The base functionality is available for all input formats compatible with the respective search engine, which Includes Thermo Fisher Scientific Orbitrap, as well as Bruker TimsTOF data.

**Figure 1.**
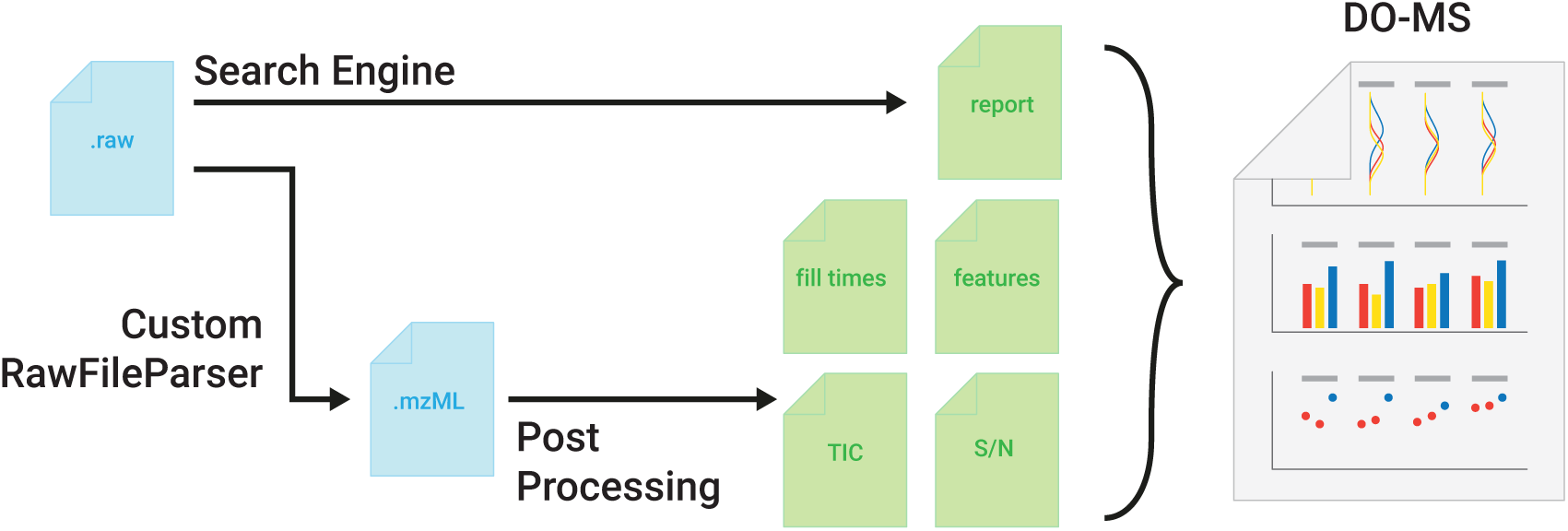
Schematic of the DO-MS pipeline version 2.0. A schematic of the processing and intermediate steps of the updated DO-MS pipeline. Input files (blue) in the raw format are searched by a search engine (the default one is DIA-NN_13_) and converted to mzML using a custom version of the ThermoRawFile parser_41_. The search report from DIA-NN and the mzML are then used by the post-processing step to analyze and display data about MS1 and MS2 accumulation times, total ion current (TIC) information, precursor-wise signal to noise levels and MS1 features.

Further, instrument-specific information is collected in a post-processing step which is only implemented for Thermo Fisher Scientific Orbitrap^42^ raw files. However, the user has the flexibility to adapt the method to other vendors, given that they can be converted to the open mzML format^43^ using tools like msConvert^44^. The current implementation uses a custom version of the ThermoRawFileParser,^41^ which reports additional instrument specific information like the noise level. It is implemented in Python^45^ and can be called from the command line which allows the search engine to automatically call post-processing after it has finished the search. General metrics like the TIC and the MS1 and MS2 accumulation times are extracted and reported in individual files. Precursor specific metrics, such as the signal to noise level (S/N), are reported based on the search engine results. Peptide like features are identified using the Dinosaur feature finder^46^. This step is independent of the amino acid sequence identification of a precursor and only based on the shape of its elution profile and isotopic envelope distribution. The metrics are then visualized in an interactive R shiny^47,48^ app, which allows the generate portable html reports. All metrics shown in this article are accessible with DO-MS and all figures resemble figures generated with DO-MS unless explicitly noted otherwise. An overview of all metric available in DO-MS can be found in the supplement *(supp. table 2)*.

### Systematic Optimization of Precursor Isolation Window Placement

In DIA experiments fragmentation spectra are highly complex due to parallel fragmentation of multiple precursors. To reduce complexity, the range of precursor masses is distributed across multiple MS2 windows, which need to be designed by the experimenter. While increasing the number of MS2 windows results in less complex spectra, it comes at the expense of an increased duty cycle length. The more MS2 scans are incorporated, the fewer data points are collected across each and every elution peak, impeding identification and optimal quantification. This trade-off needs to be optimized in a context-specific manner, depending on the sample complexity, abundance, choice of chromatography and gradient length.

DO-MS helps optimize this trade-off by systematically assessing the impact of different parameters with respect to multiple performance metrics at the same time. This is exemplified for a plexDIA experiment consisting of a 3-plex bulk lysate diluted down to the single cell level, Fig. 2. The fastest duty cycle with a single MS1 and two MS2 scans has a duration of approximately 0.9 seconds, which allows for frequent sampling of the elution profile. This results in a higher chance to sample the elution apex and is reflected in the increased MS1 peak height compared to methods with more MS2 windows, Fig. 2A,B. An acquisition method with 16 MS2 scans samples precursors only every 5.1 seconds, and thus may fail to sample the elution peak apex (*supp. table 1*). This becomes evident when the intensity of the same peptide is compared across runs. The median ratio between shared peptides is more than two-fold lower for a method with more than 12 MS2 windows compared to 2 MS2 windows, Fig. 2B. In contrast, optimal sampling of the elution apex requires more frequent sampling, which comes at the cost of fewer MS2 isolation windows. Indeed, sampling the most intense precursor signal is achieved in our experiment when using only 2 isolation windows. At the same time, such acquisition method distributes fragment ions across only two isolation windows, resulting in high co-isolation, and reduced proteome coverage. DO-MS allows to systematically and comprehensively explore this inherent trade-off between proteome coverage and sampling elution peak apexes.

**Figure 2.**
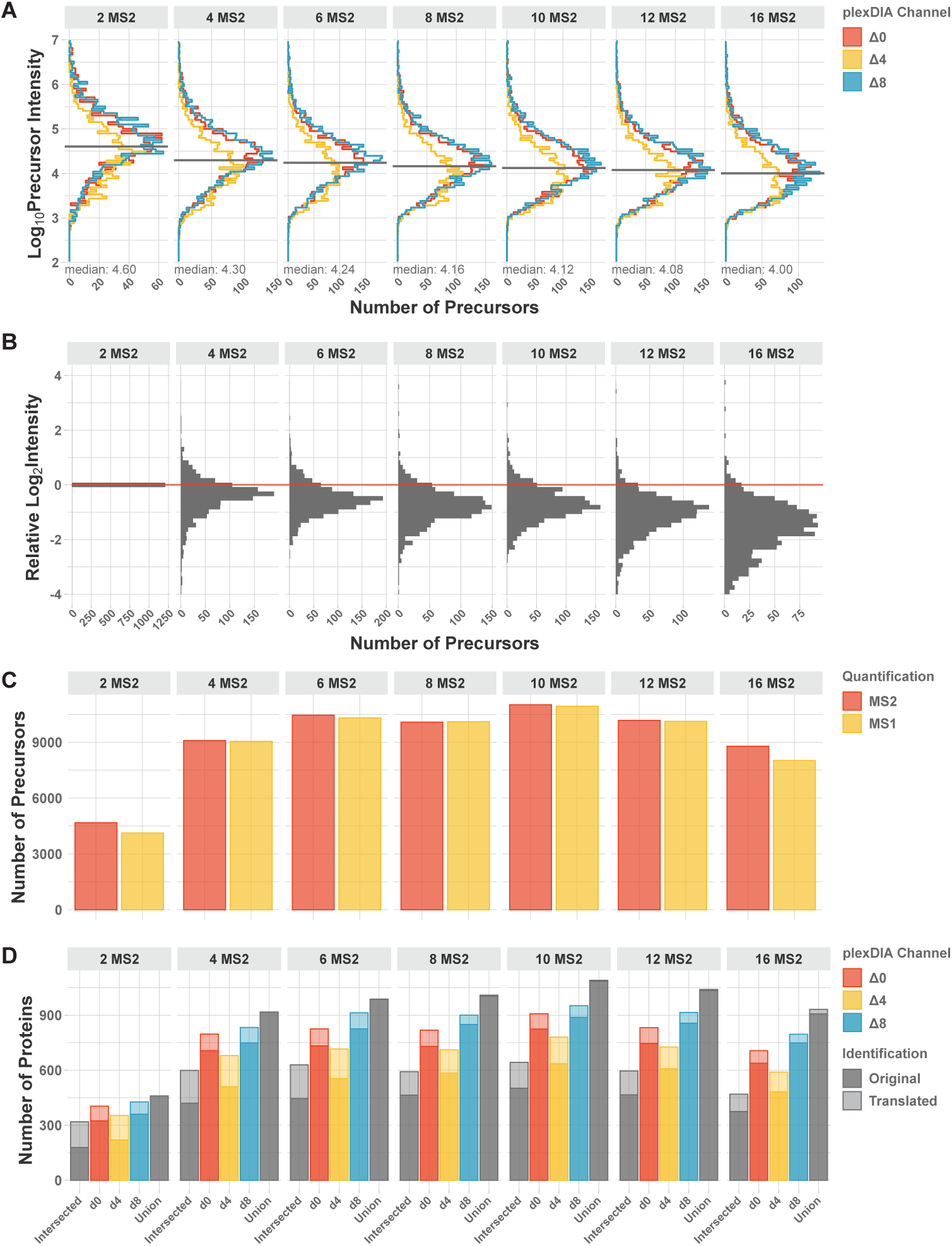
Optimizing the number of MS2 windows in the duty cycle of plexDIA methods. Example DO-MS output for a plexDIA experiment using 3-plex bulk lysate diluted down to the single-cell level with different numbers of MS2 windows. All intensities were extracted as peak heights. **A** Histogram of precursor (MS1) intensities for each plexDIA channel shown separately. **B** Distributions of ratios between precursor intensities for precursors identified across all conditions. All ratios are displayed on Log2 scale relative to the first condition. **C** The Total number of identified precursors per run is shown. Numbers are shown for precursors with MS1 (yellow) and MS2 (red) level quantification. **D** The number of protein identifications in a plexDIA set is shown for each non-isobarically labeled sample (channel). Proteins shared across all three sets and the entirety of all proteins across sets is shown in grey. Identifications which were propagated within the set are highlighted with lighter colours.

For the chosen chromatography and specimen, the DO-MS report indicates that the largest number of precursors is identified with an acquisition method of 6, 8 or 10 MS2 windows, Fig. 2C. Across all three channels about 10,000 precursors are identified on the MS2 level and quantified on the MS1 level. As we required MS2 information for sequence identification, our identifications did not benefit from the higher temporal resolution of MS1 scans and this identifications cannot exceed the number of MS2 identifications. The results indicate that overall performance balancing quantification and coverage depth is best when using 4 or 6 MS2 scans, Fig. 2. This trade-off may be mitigated by using multiple MS1 scans per duty cycle^26,27^, and such methods optimized by DO-MS using the metrics displayed in Fig. 2.

### Data Driven Optimization of Window Placement

DO-MS also allows for refinement of the precursor isolation window placement, Fig. 3. The MS2 windows can be selected to utilize equal m/z ranges^49^ or to optimize the distribution of ions across MS2 windows and thereby increase the proteome coverage^18,50^. Recently, even dynamic on-line optimization has been proposed^51^. The metrics provided by DO-MS allow users to implement previously suggested strategies or develop new ones and to continuously monitor the performance, including metrics which are often not easily accessible.

**Figure 3.**
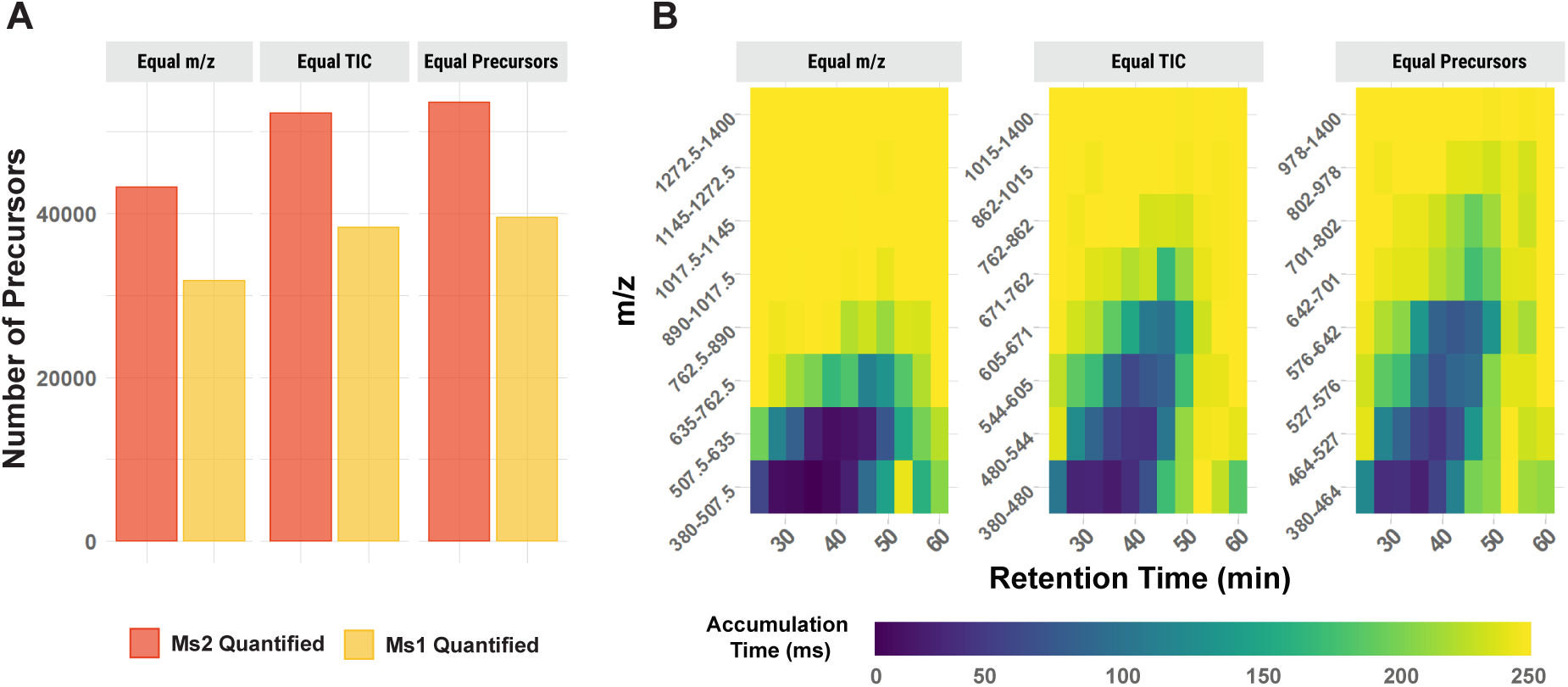
Optimizing MS2 window placement. A 3-plex experiment of 100 cell equivalent bulk lysate was analysed with 8 MS2 windows whose ranges were chosen to achieve equal distribution of (i) m/z range, (ii) ion current per window or (iii) number of precursors. **A** The total number of precursors identified on the MS2 level and quantified on the MS1 level is shown for the three different strategies. **B** The average MS2 accumulation time is shown for every MS2 window across the retention time.

As the distribution of peptide masses is not uniform across the m/z range, equal-sized isolation windows will result in more precursors per window in the lower m/z range. Thus, placement of isolation window across an equal m/z range is likely suboptimal, as manifested by lower proteome coverage shown in Fig. 3A. One of the reason for this is the associated suboptimal MS2 accumulation time, which is limited by the capacity of the ion trap. When analysing a 3-plex experiment of 100 cell equivalent bulk lysate, the lowest m/z windows will fill up in a few milliseconds, while windows with higher m/z will accumulate ions for the maximum accumulation time of 251 ms, Fig. 3B. This leads to complex fragmentation spectra, loss in sensitivity in lower mass ranges and unused ion capacity in higher m/z ranges. The effect of accumulation times on the sensitivity is likewise reflected in the lower coverage of the proteome at the MS1than at the MS2-level. The wider isolation windows at the MS1 level leads to shorter accumulation before the maximum ion trap capacity is reached. This limits sensitivity and leads to fewer quantified peptides at the MS1 than MS2 level (See also supplementary full DO-MS Report).

Windows placed based on an equal total ion current (TIC) per window, determined in a previous experiment, or based on the precursor m/z can lead to improved proteome coverage. The metrics available in DO-MS, such as accumulation times, data completeness and number of identifications as a function of FDR, allow evaluating different choices of window placement, detecting bottlenecks and improving them.

### Optimizing Chromatographic Profile and Length

To reduce the complexity of peptide sample mixtures, dimensions of separation including liquid chromatography or gas phase fractionation like trapped ion mobility spectrometry are used. Separation by liquid chromatography has been the default separation method for MS proteomics. The improved separation with longer gradients comes at the cost of increased measurement time. DO-MS allows to balance this trade-off and to perform routine quality control on peptide separation.

Longer LC gradients improve proteome coverage in DIA in two different ways. First, longer gradients lead to better separation of different peptide species reducing coelution of interferring species and improving spectral quality. Second, it leads to elongation of elution profiles resulting in precursors being sampled for a longer duration. This allows to sample each ion species less frequently and gives room for more specific isolation, improving spectral quality. Thereby, while identifying fewer peptides per unit time, longer gradients facilitate identifying more peptides per sample. The general trend is shown by the DO-MS output for a 3-plex 100-cell equivalent bulk dilution analyzed with 15, 30 and 60 minutes of active gradient using the same duty cycle, Fig. 4. One benefit of the longer gradients can be seen when the ion accumulation time of the Orbitrap instrument is plotted as a function of the retention time, Fig. 4A. Longer gradients distribute the analytes and lead to longer accumulation of ions, before the maximum capacity is reached. Individual spectra therefore contain fewer ion species and sample sufficient ions even from low abundant peptides. This improves not only the absolute numbers of identifications but also the fraction of precursors quantified at the MS1-level, Fig. 4B.

**Figure 4.**
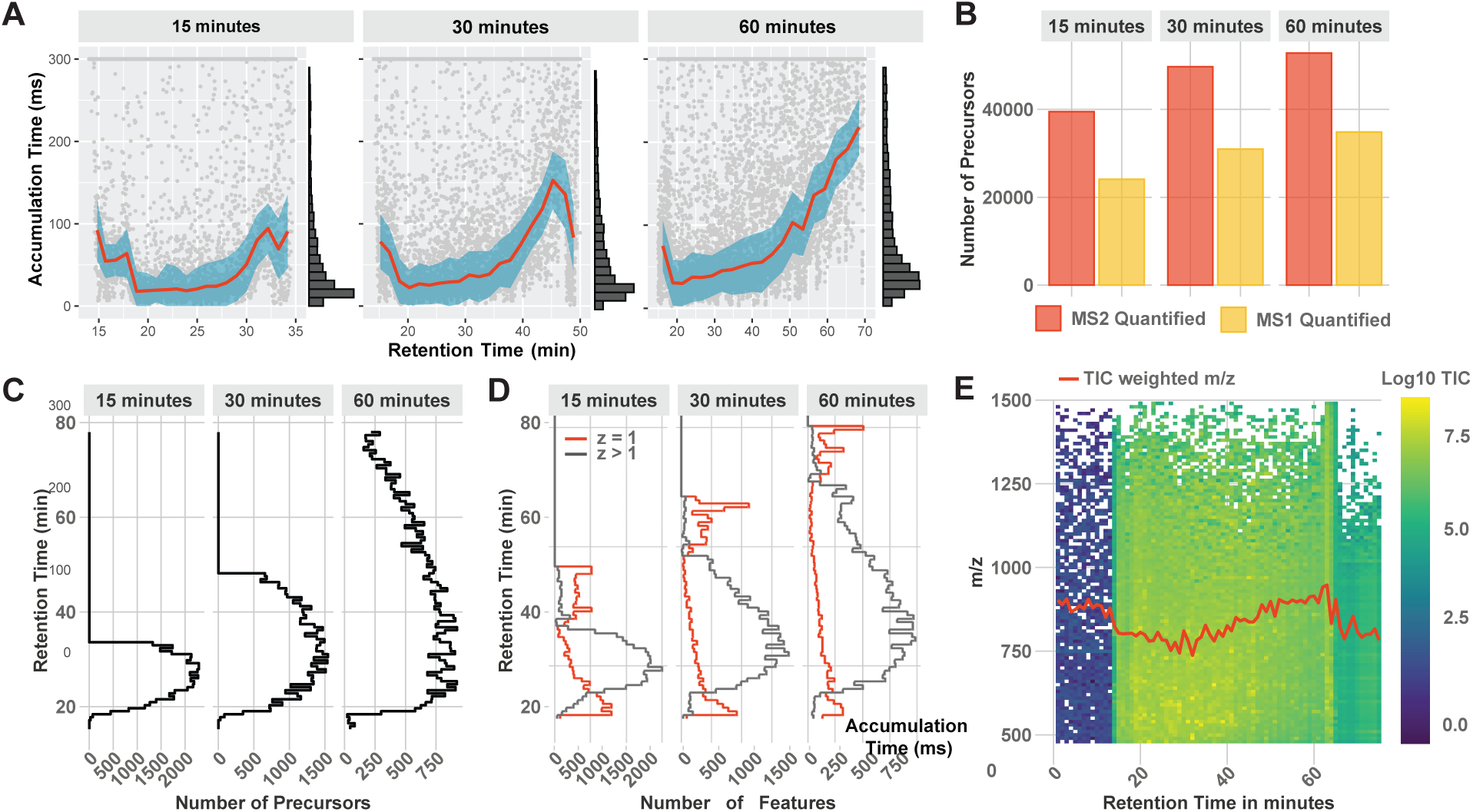
Optimizing gradient profile and length. DO-MS allows to optimize the LC gradient of experiments based on metrics capturing the whole LC-MS workflow. **A** The distribution of MS1 accumulation times across the LC gradient. **B** Number of quantified precursors in relation to the gradient length. **C** Number of identified precursors by the search engine across the gradients and **D** ion features identified by Dinosaur. **E** Ion map displaying the TIC and mean m/z (red curve) as a function of the retention time. All data are from 100x 3-plexDIA samples as described in the methods.

DO-MS also allows to optimize the slope and profile of the gradient to evenly distribute ions across a gradient while keeping its duration constant. Depending on the sample, peptides might not elute evenly across the gradient. This information becomes accessible in three different ways. DO-MS reports the accumulation time of the ion trap Fig. 4A, peptide identifications across the gradient Fig. 4C, and peptide like features or potential contaminants assembled by Dinosaur across the gradient Fig. 4D.

Having access to gradient specific parameters facilitates effective quality control and problem identification. Identified MS1 features provide useful information for ion clusters not assigned to a peptide sequence including singly charged species and peptide-like ions not mapped to a sequence, Fig. 4D. This can be useful to identify contaminants^31^ and estimate the ions accessible to MS analysis that may be interpreted by improved algorithms^8,52^. The binned TIC output allows to identify errors in the method setup and gives a quick overview of the sampled mass range, Fig. 4E.

### Improving Sampling Using Additional Survey Scans

The conflict between reducing spectral complexity and increasing the number of data points per peak mentioned in Fig. 2 can be partially alleviated by increasing the number of survey scans^27^. When duty cycles are long, more frequent sampling on the MS1 level can increase the fraction of precursors with MS1 information and the probability of sampling close to the elution apex^19,26^. The DO-MS framework can be used to assess the contribution of such additional MS1 scans to improved precursor sampling.

The effect can be exemplified based on 3-plexDIA set whose samples correspond to 100-cells per channel analyzed analyzed with 60 minutes of active gradient. A method with a single survey scan is compared to a method with two survey scans evenly distributed between the eight MS2 scans, Fig. 5A. The additional survey scan increases the duty cycle length only marginally, while increasing the frequency of precursor sampling almost 2-fold. Thus, the adapted method increases the probability that precursors are sampled close to their elution apex and that peptides with a shorter elution profile and potentially lower intensity can be quantified on the MS1 level, which would be otherwise missed. These expectations are supported by the results shown in Fig. 5B-D.

**Figure 5.**
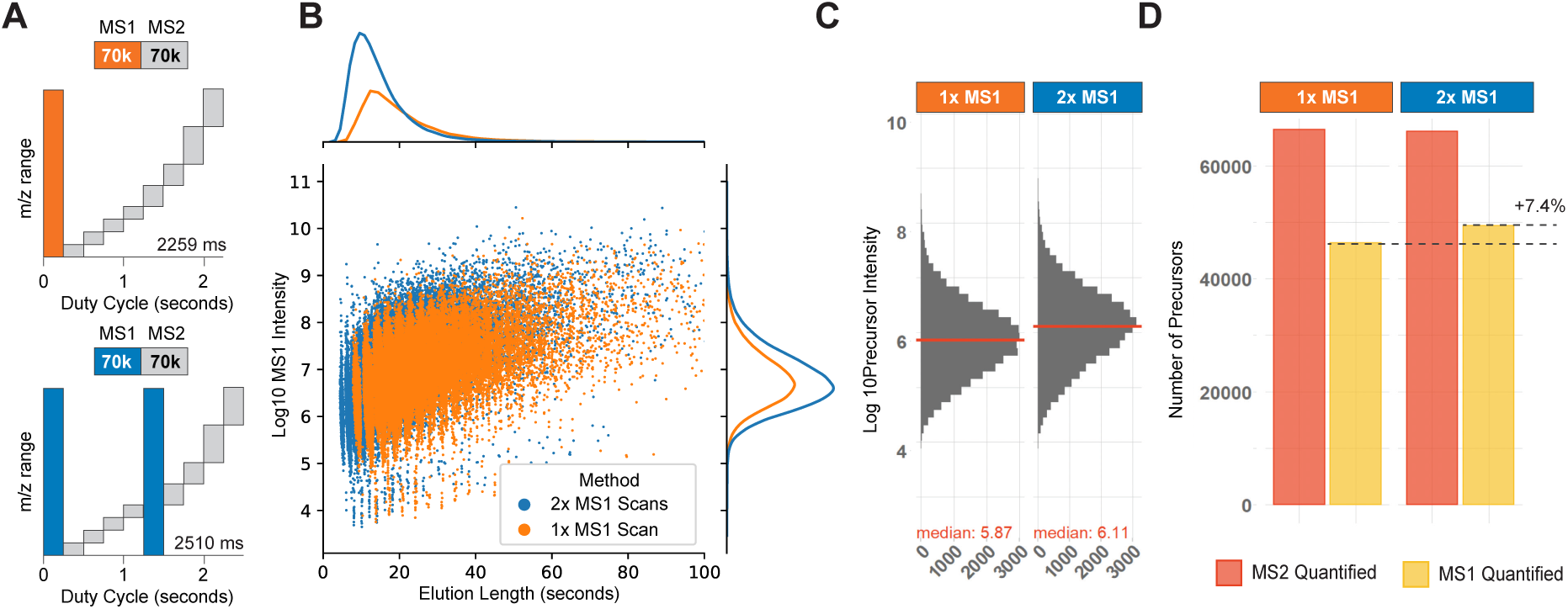
Effect of additional survey scans per duty cycle. Data acquisition methods can employ multiple survey scans to improve precursor sampling and reduce stochastic sampling effect. **A** Diagrams of a duty cycle with a single survey scan (orange) and a duty cycle with two survey scans (blue). **B** All peptide like features identified by Dinosaur_46_ are displayed with their elution length at base and MS1 intensity. The associated marginal distributions are shown. The additional survey scan allows to detect many additional peptide-like features with shorter elution profile. **C** The MS1 intensity of intersected precursors is increased upon introduction of an additional survey scan. **D** The fraction of MS1 quantified precursors is increased with additional survey scans while maintaining the total number of identifications, independent of the slightly increased duty cycle time. The data shows a 100-cell equivalent 3-plex dataset acquired on 60 min active gradient as described in the methods. Panel B was plotted outside of DO-MS using the peptide-like feature information as stated in the methods.

More survey scans lead to almost doubling the number of identified peptide-like features, with the increase being particularly pronounced for features with short elution length, Fig. 5B. The improvements also result higher MS1 intensity estimates by the search engine for intersected precursors since more precursors are samples close to their apexes. Furthermore, a larger fraction of precursors are quantified at the MS1, Fig. 5C,D. These improvements are observed without associated negative effects due to the longer overall duty cycle. These results indicate that the duty cycle with 2 MS1 survey scans outperforms the one with single MS1 survey scans.

### Quality Control for Routine Sample Acquisition

When acquiring large datasets, it is important to continuously monitor the performance of the acquisition method and identify potential failed experiments^37^. This monitoring for plexDIA experiments should include metrics for each labeled sample i.e., channel level metrics.

DO-MS provides a convenient way to perform such quality control, exemplified by the singlecell plexDIA set by Derks *et al.*^26^ shown in Fig. 6. Using nPOP sample preparation^53^, 10 sets with 3 single cells each were prepared and measured on a timsTOF instrument, resulting in about 1,000 quantified proteins per single cell on average, Fig. 6A. As plexDIA can benefit from translating precursor identifications between channels^26,27^, the impact of translation on identifications and data completeness is reported by DO-MS. With single cells it is vital to identify potential dropouts where sample preparation might have failed and exclude them from processing. One useful metric for this is the precursor intensity distribution for every single cell, which is displayed by DO-MS, Fig. 6B. Another metric to assess the single-cell proteome quality is the quantification variability between peptides originating from the same protein, which has been proposed as a metric for single-proteome quality^54^, Fig. 6C. In this dataset, the cells in channel Δ0, set 06 and Δ8, set 10 show both lower number of proteins before translation and a higher quantification variability, and should potentially be excluded from further analysis.

**Figure 6.**
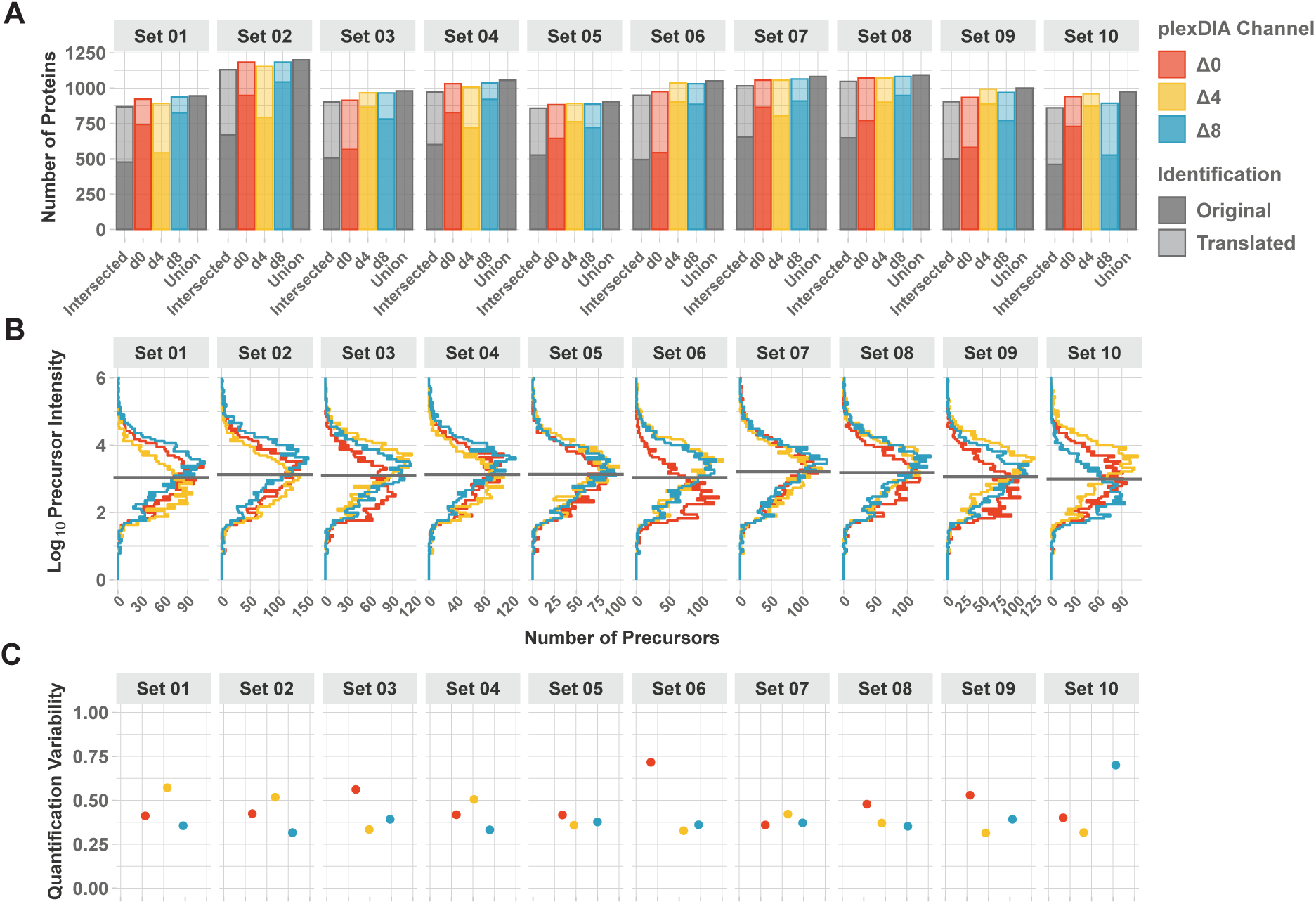
Routine quality control. When acquiring data from a large number of single cells, DO-MS can be used to get a quick overview of the quality of the processing results. **A** Number of protein identifications per single cell before and after translating identifications between channels. Only identifications quantified on the MS1 level are shown. **B** Channel wise intensity distribution of identified precursors. **C** Quantification variability calculated as the coefficient of variation between peptides of the same protein. The report was generated from the data published by Derks *et al.*^26^ for 10 single-cell 3-plex sets analyzed on a timsTOF instrument.

### Conclusion

The DO-MS framework provides a systematic approach to benchmarking, optimizing, and reporting results from label free and multiplexed DIA-MS. We exemplified how key method parameters such as the number of precursor scans or isolation window placement can be benchmarked and optimized. DO-MS aims to foster understanding from first-principles, considering fundamental trade-offs such as spectral complexity and sampling frequency. By adopting this approach, it becomes possible to design methods tailored to specific application needs, such as emphasizing data completeness, quantitative accuracy, or proteome depth. DO-MS should enable broader adoption of cutting edge methods DIA and plexDIA methods for driving biological research^55^.

## Methods

### Data Acquisition

Apart from the 30 single cells acquired on the timsTOF as part of plexDIA, all samples consist of bulk cellular lysates diluted down to the respective number of single-cell equivalents by assuming a 250pg of protein per cell. Melanoma cells (WM989-A6-G3, a kind gift from Arjun Raj, University of Pennsylvania), U-937 cells (monocytes), and HPAF-II cells (PDACs, American Type Culture Collection (ATCC), CRL-1997) were cultured as previously described by Derks *et al.*^26^ – methods - cell culture. Cells were harvested, processed, and labeled with mTRAQ as described by Derks *et al.*^26^ - methods - Preparation of bulk plexDIA samples.

All bulk data was acquired on the Thermo Fisher Scientific Q-Exactive Classic Orbitrap mass spectrometer. Samples consisting of 1-*µ*l were injected with the Dionex UltiMate 3000 UHPLC using 25 cm×75µm IonOpticks Aurora Series UHPLC column (AUR2-25075C18A). Two buffers A and B were used with buffer A made of 0.1% formic acid (Pierce, 85178) in LC–MS-grade water and buffer B made of 80% acetonitrile and 0.1% formic acid mixed with LC–MS-grade water.

### Systematic optimization of precursor isolation windows

A combined sample consisting of one single-cell equivalent PDAC lysate labeled with mTRAQd0, one single-cell equivalent U937 lysate labeled with mTRAQd4 and one single-cell equivalent Melanoma lysate labeled with mTRAQd8 was injected with 1ul volume. Liquid chromatography was performed with 200nl/min for 30 minutes of active gradient starting with 4% Buffer B (minutes 0–2.5), 4–8% B (minutes 2.5–3), 8–32% B (minutes 3–33), 32–95% B (minutes 33–34), 95% B (minutes 34–35), 95-4% B (minutes 35–35.1), 4% B (minutes 35.1-53). All acquisition methods had a single MS1 scan covering the range of 380mz-1400mz followed by DIA MS2 scans: 2xMS2 starting at 380mz: 240Th, 780Th width; 4xMS2 starting at 380mz: 120Th, 120Th, 200Th, 580Th width; 6xMS2 starting at 380mz: 80Th, 80Th, 80Th, 120Th, 240Th, 420Th width; 8xMS2 starting at 380mz: 60Th, 60Th, 60Th, 60Th, 100Th, 100Th, 290Th, 290Th width; 10xMS2 starting at 380mz: 50Th, 50Th, 50Th, 50Th, 50Th, 75Th, 75Th, 150Th, 150Th, 320Th width; 12xMS2 starting at 380mz: 40Th, 40Th, 40Th, 40Th, 40Th, 40Th, 60Th, 60Th, 120Th, 120Th, 210Th, 210Th width; 16xMS2 starting at 380mz: 30Th, 30Th, 30Th, 30Th, 30Th, 30Th, 30Th, 30Th, 50Th, 50Th, 50Th, 50Th, 145Th, 145Th, 145Th, 145Th width. All MS1 and MS2 scans were performed with 70,000 resolving power, 3×10^6^ AGC maximum, 300-ms maximum accumulation time, NCE at 27%, default charge of 2, and RF S-lens was at 80%.

### Data Driven optimization of window placement

A combined sample consisting of 100 single-cell equivalents of PDAC, U937, and Melanoma cells were labled with mTRAQd0, mTRAQd4 and mTRAQd8 respectively. Liquid chromatography was performed with 200nl/min for 30 minutes of active gradient starting with 4% Buffer B (minutes 0–2.5), 4–8% B (minutes 2.5–3), 8–32% B (minutes 3–33), 32–95% B (minutes 33–34), 95% B (minutes 34–35), 95-4% B (minutes 35–35.1), 4% B (minutes 35.1-53). Both MS1 and MS2 scans covered a range of 380mz to 1400mz with a single MS1 scan and 8 MS2 scans. The distribution of precursors was determined based on DO-MS report using equal sized windows, starting at 380mz: 127.5Th, 127.5Th, 127.5Th, 127.5Th, 127.5Th, 127.5Th, 127.5Th, 127.5Th width. MS2 windows where then distributed to have equal TIC based on the DO-MS output: starting at 380mz: 100Th, 64Th, 61Th, 66Th, 91Th, 100Th, 153Th, 385Th. For the equal number of precursors, the original sample was searched with DIANN as described and MS2 windows were distributed to have an equal number of precursors: starting at 380mz: 84Th, 63Th, 49Th, 66Th, 59Th, 101Th, 176Th, 422Th. All MS1 and MS2 scans were performed with 70,000 resolving power, 3×10^6^ AGC maximum, 251-ms maximum accumulation time, NCE at 27%, default charge of 2, and RF S-lens was at 80%.

### Optimizing gradient profile and length

A combined sample consisting of 100 single-cell equivalents of PDAC, Melanoma and U937 were labled with mTRAQd0, mTRAQd4 and mTRAQd8 respectively. Liquid chromatography was performed with 200nl/min flow rate starting with 4% Buffer B (minutes 0–2.5) followed by 4–8% B (minutes 2.5–3). The active gradient with 8% buffer B to 32% buffer B stretched across 15, 30 and 60 minutes followed by a 1 minute 32–95% B ramp, 1 minute at 95% and 18 minutes at 4% B. All acquisition methods had a single MS1 scan covering the range of 478mz-1500mz followed by 8 DIA MS2 scans: starting at 380mz: 60Th, 60Th, 60Th, 60Th, 100Th, 100Th, 290Th, 290Th. All MS1 and MS2 scans were performed with 70,000 resolving power, 3×10^6^ AGC maximum, 300-ms maximum accumulation time, NCE at 27%, default charge of 2, and RF S-lens was at 80%.

### Effect of additional survey scans

A 100 single-cell equivalent of each, PDAC, U937 and Melanoma cells were labeled with mTRAQd0, mTRAQd4 and mTRAQd8 respectively and injected in a volume of 1ul. Liquid chromatography was performed with 200nl/min for 30 minutes of active gradient starting with 4% Buffer B (minutes 0–2.5), 4–8% B (minutes 2.5–3), 8–32% B (minutes 3–63), 32–95% B (minutes 63–64), 95% B (minutes 64–65), 95-4% B (minutes 65–65.1), 4% B (minutes 65.1-83). A single MS1 scan with a range of 478mz-1500mz was followed by MS2 scans starting at 380mz with 60Th, 60Th, 60Th, 60Th, 100Th, 100Th, 290Th, 290Th width. For the method with increased MS1 sampling, a second MS1 scan was incorporated after the fourth MS2 scan. All MS1 and MS2 scans were performed with 70,000 resolving power, 3×10^6^ AGC maximum, 251-ms maximum accumulation time, NCE at 27%, default charge of 2, and RF S-lens was at 80%.

### Data Analysis

Data was analysed using DIA-NN 1.8.1 using the 5,000 protein group human-only spectral library published previously by Derks *et al.*^26^ - methods - Spectral library generation. Data was then processed with DO-MS. For preprocessing of Orbitrap data DO-MS uses ThermoRawFileParser 1.4.0 to convert the proprietary raw format to the open mzML standard and Dinosaur 1.2.0 for feature detection. All other preprocessing steps are performed in the Python programming language version 3.10 and makes use of its extensive ecosystem for scientific programming including Numpy, Pandas, pymzML and scikit-learn. All plots were created in DO-MS which utilizes the R programming language version 4.3.1. Figure 5B was created using matplotlib.

Data completeness is shown for all pairwise comparisons in a plex DIA set. It is calculated as the Jaccard index between two sets of identifications *A* and *B* given by:

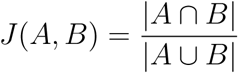

## Availability

Further documentation on the use of DO-MS is available at do-ms.slavovlab.net. The current version 2.0 is open source and freely available at github.com/SlavovLab/DO-MS. All data shown as example application is available at do-ms.slavovlab.net/docs/DO-MS_examples. The 30 single cells plexDIA dataset acquired on the timsTOF has been published as part of plexDIA and is available at http://scp.slavovlab.net/Derks_et_al_2022. blue All other data acquired for this study has been deposited on MassIVE under the accession MSV000091733.

## Acknowledgments

We thank Luke Khoury for support with sample processing and acquisition and Jason Derks for sample preparation. The work was funded by an Allen Distinguished Investigator award through The Paul G. Allen Frontiers Group to N.S., a Seed Networks Award from CZI CZF2019-002424 to N.S., an NIGMS award R01GM144967 to N.S., and an NCI award UG3CA268117 to N.S.

