## Supplemental table and DO-MS reports for "Data-Driven Optimization of DIA Mass Spectrometry by DO-MS": 01_DO-MS_Report_MS1_number.html (from another version of this preprint: version 2, 2023-07-31)


### DO-MS Report

###### Version: 2.0.6

#### Summary

##### DIA-NN Experiments

DIA-NN Experiments

2 MS2 F:/GW/raw\_data/wGW026.raw

4 MS2 F:/GW/raw\_data/wGW027.raw

6 MS2 F:/GW/raw\_data/wGW025.raw

8 MS2 F:/GW/raw\_data/wGW028.raw

10 MS2 F:/GW/raw\_data/wGW029.raw

12 MS2 F:/GW/raw\_data/wGW030.raw

16 MS2 F:/GW/raw\_data/wGW031.raw

#### Ion Sampling

##### Channel wise MS1 Intensity for Precursors

Plotting the MS1 intensity for all precursors which were associated
with one of the defined channels.

##### Precursors Identified across Gradient

Precursor are plotted across the chromatographic gradient.

##### MS1 Intensity summed over all Channels.

Plotting the MS1 intensity for all precursors summed over all
channels.

##### MS1 Intensity for Intersected Precursors, summed over all Channels.

Plotting the MS1 intensity for all precursors summed over all
channels. Only intersected precursors present in all loaded experiments
are shown.

##### Normalized MS1 Intensity for Intersected Precursors

Plotting the MS1 Intensity for intersected precursors summed over all
channels. Experiments are normalized to the first experiment.

##### Number of Precursors by Charge State

Number of precursors observed during MS1 scans by charge state

##### Ms1 Fill Time Distribution

Ms1 fill times along gradient

##### Ms1 Fill Times along Gradient

The averge fill time is shown in magenta for different bins along the
retention time gradient. The standard deviation is depicted as area in
blue, scans outside this area are shown as single datapoints.

##### Ms1 total Ion Current along Gradient

The toal Ion Current (TIC) is shown for bins along the retention time
gradient.

```
## [1] "Plot failed to render. Reason: Error: Upload tic.tsv\n"
```

##### Ms2 Fill Time Distribution

Ms2 fill times along gradient

##### Ms2 Fill Times along Gradient

The averge fill time is shown in magenta for different bins along the
retention time gradient. The standard deviation is depicted as area in
blue, scans outside this area are shown as single datapoints.

##### Ms2 Fill Time Matrix

The average Ms2 fill times are shown across the gradient for every
distinct Ms2 window.

```
## [1] "Plot failed to render. Reason: Error: Your fill_times.tsv file does not contain the Window.Lower column for the isolation window. Please make sure that you use the latest version of DO-MS DIA.\n"
```

##### Channel wise MS1 Copy Number for Precursors

Plotting the MS1 copy numbers for all precursors which were
associated with one of the defined channels. The copy numbers are
calculated using the signal to noise ratio as described in Derks et
al. 2022. By default a resolution of 70,000 is used during
preprocessing. It can be changed with the –resolution parameter

```
## [1] "Plot failed to render. Reason: Error: Upload report.txt\n"
```

#### Identifications

##### Number of Confident Precursor Identifications

Plotting the number of precursors identified at each given confidence
level.

##### Precursors by Quantification Strategy

Number of precursors found based on quantification startegy. Ms2
precursors are counted based on Precursor.Quantity > 0 and Ms1
precursors are counted based on Ms1.Area > 0.

##### Precursors by Modification

Number of precursors found based on modification types specified

##### Miscleavage Rate (percentage), PEP < 0.01

Miscleavage rate (percentage) for precursors identified with
confidence PEP < 0.01

##### Miscleavage Rate (K), PEP < 0.01

Plotting frequency of lysine miscleavages in confidently identified
precursors.

##### Miscleavage Rate (R), PEP < 0.01

Plotting frequency of arginine miscleavages in confidently identified
precursors.

##### Number of Protein Identifications

Number of proteotypic protein IDs found per run. Protein IDs are
shown across all channels in an experiment.

#### plex-DIA Diagnostics

##### Identified Precursors per Channel

The number of precursors identified is shown for every channel
together with the number of total and intersected precursors. The number
of precursors is based on all precursors found in the report.tsv file
which is by default controlled by a run-specific FDR.

##### Identified Precursors per Channel, Channel Q-Value

The number of precursors identified is shown for every channel
together with the number of total and intersected precursors. The number
of precursors is based on all precursors with a Channel.Q.Value <=
0.01.

##### Identified Proteins per Channel

The number of Proteins identified is shown for every channel together
with the number of total and intersected proteins. The number of
proteins is based on all proteotypic precursors independent of the
Protein.Q.Value.

##### Relative Single-Cell Intensity

Single-Cell intensity relative to the carrier channel for intersected
precursors

##### MS1 Quantification Variability

Single-Cell intensity relative to the carrier channel for intersected
precursors

##### MS2 Quantification Variability

Single-Cell intensity relative to the carrier channel for intersected
precursors

##### Total MS1 Precursors

Total number of precursors identified based on different confidence
metrics.

```
## [1] "Plot failed to render. Reason: Error: Upload ms1_extracted.tsv\n"
```

##### Missing Data, Precursor Level

Plotting the Jaccard Index for identified precursors for all channel
combinations.

##### Missing Data, Protein Level

Plotting the Jaccard Index for identified precursors for all channel
combinations.

#### Feature Detection

##### Features Identified by Charge

Identified features are reported based on the charge. Precursors
quantified in seperate channels are treated as separate precursors..

##### Isotopic Peaks Identified per Feature

The number of isotopic peaks identified is shown for features
detected in the Dinosaur search.

##### Number of Scans per feature

The number of MS1 scans is shown for all features identified in the
Dinosaur search.

##### Retention Length of Features at Base

Plotting the retention length of identified features at the base.

##### Features Identified across Gradient

The frequency of precursor identifications based on the Dinosaur
search is plotted across the chromatographic gradient.

##### Features Identified across m/z

The frequency of precursor identifications based on the Dinosaur
search is plotted across the mass to charge ratio.

##### Feature Intensity Distribution

The distribution of integrated intensities is shown for identified
features.
