## Supplemental table and DO-MS reports for "Data-Driven Optimization of DIA Mass Spectrometry by DO-MS": 02_DO-MS_Report_MS2_strategies.html (from another version of this preprint: version 2, 2023-07-31)


### DO-MS Report

###### Version: 2.0.6

#### Summary

##### DIA-NN Experiments

DIA-NN Experiments

Equal m/z F:/GW/raw\_data/wGW061.raw

Equal TIC F:/GW/raw\_data/wGW062.raw

Equal Precursor F:/GW/raw\_data/wGW063.raw

#### Ion Sampling
