## Supplemental table and DO-MS reports for "Data-Driven Optimization of DIA Mass Spectrometry by DO-MS": 03_DO-MS_Report_gradient_length.html (from another version of this preprint: version 2, 2023-07-31)


### DO-MS Report

###### Version: 2.0.6

#### Summary

##### DIA-NN Experiments

DIA-NN Experiments

15 minutes F:/GW/raw\_data/eGW053.raw

30 minutes F:/GW/raw\_data/eGW054.raw

60 minutes F:/GW/raw\_data/eGW056.raw

#### Ion Sampling

##### Channel wise MS1 Intensity for Precursors
