## Supplemental table and DO-MS reports for "Data-Driven Optimization of DIA Mass Spectrometry by DO-MS": 05_DO-MS_Report_sc_qc.html (from another version of this preprint: version 2, 2023-07-31)


### DO-MS Report

###### Version: 2.0.6

#### Summary

##### DIA-NN Experiments

DIA-NN Experiments

Set 1
F:/JD/plexDIA/Bruker/OneDrive\_1\_3-9-2022/20220308\_npop\_SCP\_1\_Slot2-1\_1\_215.d

Set 2
F:/JD/plexDIA/Bruker/OneDrive\_1\_3-9-2022/20220308\_npop\_SCP\_2\_Slot2-2\_1\_216.d

Set 3
F:/JD/plexDIA/Bruker/OneDrive\_1\_3-9-2022/20220308\_npop\_SCP\_3\_Slot2-3\_1\_217.d

Set 4
F:/JD/plexDIA/Bruker/OneDrive\_1\_3-9-2022/20220308\_npop\_SCP\_4\_Slot2-4\_1\_218.d

Set 5
F:/JD/plexDIA/Bruker/OneDrive\_1\_3-9-2022/20220308\_npop\_SCP\_5\_Slot2-5\_1\_219.d

Set 6
F:/JD/plexDIA/Bruker/OneDrive\_1\_3-9-2022/20220308\_npop\_SCP\_6\_Slot2-6\_1\_220.d

Set 7
F:/JD/plexDIA/Bruker/OneDrive\_1\_3-9-2022/20220308\_npop\_SCP\_7\_Slot2-7\_1\_221.d

Set 8
F:/JD/plexDIA/Bruker/OneDrive\_1\_3-9-2022/20220308\_npop\_SCP\_8\_Slot2-8\_1\_222.d

Set 9
F:/JD/plexDIA/Bruker/OneDrive\_1\_3-9-2022/20220308\_npop\_SCP\_9\_Slot2-9\_1\_223.d

Set 10
F:/JD/plexDIA/Bruker/OneDrive\_1\_3-9-2022/20220308\_npop\_SCP\_10\_Slot2-10\_1\_224.d

#### Ion Sampling

##### Channel wise MS1 Intensity for Precursors

Plotting the MS1 intensity for all precursors which were associated
with one of the defined channels.

##### Precursors Identified across Gradient

Precursor are plotted across the chromatographic gradient.

##### MS1 Intensity summed over all Channels.

Plotting the MS1 intensity for all precursors summed over all
channels.

##### MS1 Intensity for Intersected Precursors, summed over all Channels.

Plotting the MS1 intensity for all precursors summed over all
channels. Only intersected precursors present in all loaded experiments
are shown.

##### Normalized MS1 Intensity for Intersected Precursors

Plotting the MS1 Intensity for intersected precursors summed over all
channels. Experiments are normalized to the first experiment.

##### Number of Precursors by Charge State

Number of precursors observed during MS1 scans by charge state

##### Ms1 Fill Time Distribution

Ms1 fill times along gradient

```
## [1] "Plot failed to render. Reason: Error: Upload fill_times.txt\n"
```

##### Ms1 Fill Times along Gradient

The averge fill time is shown in magenta for different bins along the
retention time gradient. The standard deviation is depicted as area in
blue, scans outside this area are shown as single datapoints.

```
## [1] "Plot failed to render. Reason: Error: Upload fill_times.txt\n"
```

##### Ms1 total Ion Current along Gradient

The toal Ion Current (TIC) is shown for bins along the retention time
gradient.

```
## [1] "Plot failed to render. Reason: Error: Upload tic.tsv\n"
```

##### Ms2 Fill Time Distribution

Ms2 fill times along gradient

```
## [1] "Plot failed to render. Reason: Error: Upload fill_times.txt\n"
```

##### Ms2 Fill Times along Gradient

The averge fill time is shown in magenta for different bins along the
retention time gradient. The standard deviation is depicted as area in
blue, scans outside this area are shown as single datapoints.

```
## [1] "Plot failed to render. Reason: Error: Upload fill_times.txt\n"
```

##### Ms2 Fill Time Matrix

The average Ms2 fill times are shown across the gradient for every
distinct Ms2 window.

```
## [1] "Plot failed to render. Reason: Error: Upload tic.tsv\n"
```

#### Feature Detection

##### Features Identified by Charge

Identified features are reported based on the charge. Precursors
quantified in seperate channels are treated as separate precursors..

```
## [1] "Plot failed to render. Reason: Error: Please provide a features.tsv file\n"
```

##### Isotopic Peaks Identified per Feature

The number of isotopic peaks identified is shown for features
detected in the Dinosaur search.

```
## [1] "Plot failed to render. Reason: Error: Please provide a features.tsv file\n"
```

##### Number of Scans per feature

The number of MS1 scans is shown for all features identified in the
Dinosaur search.

```
## [1] "Plot failed to render. Reason: Error: Please provide a features.tsv file\n"
```

##### Retention Length of Features at Base

Plotting the retention length of identified features at the base.

```
## [1] "Plot failed to render. Reason: Error: Please provide a features.tsv file\n"
```

##### Features Identified across Gradient

The frequency of precursor identifications based on the Dinosaur
search is plotted across the chromatographic gradient.

```
## [1] "Plot failed to render. Reason: Error: Please provide a features.tsv file\n"
```

##### Features Identified across m/z

The frequency of precursor identifications based on the Dinosaur
search is plotted across the mass to charge ratio.

```
## [1] "Plot failed to render. Reason: Error: Please provide a features.tsv file\n"
```

##### Feature Intensity Distribution

The distribution of integrated intensities is shown for identified
features.

```
## [1] "Plot failed to render. Reason: Error: Please provide a features.tsv file\n"
```
